# A replicon RNA vaccine induces durable protective immunity from SARS-CoV-2 in nonhuman primates after neutralizing antibodies have waned

**DOI:** 10.1101/2022.08.08.503239

**Authors:** Megan A. O’Connor, David W. Hawman, Kimberly Meade-White, Shanna Leventhal, Wenjun Song, Samantha Randall, Jacob Archer, Thomas B. Lewis, Brieann Brown, Naoto Iwayama, Chul Ahrens, William Garrison, Solomon Wangari, Kathryn A. Guerriero, Patrick Hanley, Jamie Lovaglio, Greg Saturday, Paul T. Edlefsen, Amit Khandhar, Heinz Feldmann, Deborah Heydenburg Fuller, Jesse H. Erasmus

## Abstract

The global SARS-CoV-2 pandemic prompted rapid development of COVID-19 vaccines. Although several vaccines have received emergency approval through various public health agencies, the SARS-CoV-2 pandemic continues. Emergent variants of concern, waning immunity in the vaccinated, evidence that vaccines may not prevent transmission and inequity in vaccine distribution have driven continued development of vaccines against SARS-CoV-2 to address these public health needs. In this report, we evaluated a novel self-amplifying replicon RNA vaccine against SARS-CoV-2 in a pigtail macaque model of COVID-19 disease. We found that this vaccine elicited strong binding and neutralizing antibody responses. While binding antibody responses were sustained, neutralizing antibody waned to undetectable levels after six months but were rapidly recalled and conferred protection from disease when the animals were challenged 7 months after vaccination as evident by reduced viral replication and pathology in the lower respiratory tract, reduced viral shedding in the nasal cavity and lower concentrations of pro-inflammatory cytokines in the lung. Cumulatively, our data demonstrate in pigtail macaques that a self-amplifying replicon RNA vaccine can elicit durable and protective immunity to SARS-CoV-2 infection. Furthermore, these data provide evidence that this vaccine can provide durable protective efficacy and reduce viral shedding even after neutralizing antibody responses have waned to undetectable levels.

## INTRODUCTION

The SARS-CoV-2 pandemic continues with resurgent case counts in multiple countries. Despite rapid development of several vaccine candidates and rollout of these vaccines in developed countries, breakthrough infections, the emergence of variants of concern (VoCs) that are more resistant to vaccine-induced immunity, and the inequity in vaccine distribution between wealthy and developing countries emphasize the need for continued development of new vaccines for SARS-CoV-2 that can address these gaps. In addition, waning immunity in vaccinated populations has become a public health concern and booster shots are now recommended to sustain immunity induced by currently licensed COVID-19 vaccines to provide cross-protection from emerging VoCs. To address the public health crisis presented by COVID-19, an ideal vaccine should reduce transmission potential by protecting the upper airway from infection, provide protection from severe respiratory disease, and elicit sustained immunity against SARS-CoV-2 without the need for frequent booster immunizations.

Although the correlates of COVID-19 vaccine protection are still under investigation, evidence for the currently licensed vaccines supports a dominant role of neutralizing antibody responses (*1–3*). Viral infection or vaccination induces rapid expansion of effector cells needed to combat an infection, yet over time, there is a natural contraction of this response, leaving a pool of memory T- and B-cells and effector plasma cells that can rapidly respond and clear a secondary infection. Neutralizing and non-neutralizing antibody responses following vaccination with the first generation two-dose COVID-19 vaccines, mRNA-1273 and BNT162b2, wane within 6 months after vaccination (*4–7*), but whether sustained levels of circulating antibody are required to maintain protection from disease is still not clear. The waves of SARS-CoV-2 infections driven by the emergence of VoCs and the waning of vaccine-induced antibody responses (*8*) have prompted recommendations for COVID-19 booster immunizations. Indeed, booster immunizations with vaccines directed towards the parental lineage enhance cross-neutralizing antibodies against VoCs, including Omicron, and contribute to reduced COVID-19 disease severity and death during breakthrough infections (*4, 9, 10*). While it is now clear that rapid waning of neutralizing antibody coupled with the emergence of more transmissible and resistant VoCs are permitting breakthrough infections after vaccination, the precise immune mechanisms contributing to the preservation of vaccine-induced protection from disease are not yet clear.

Previously, we reported that an alphavirus-derived replicon RNA (repRNA) SARS-CoV-2 vaccine encoding the SARS-CoV-2 spike protein and formulated with a novel cationic nanocarrier, that offers the advantages of superior stability at warmer temperatures and rapid scale-up (repRNA-CoV2S), induced robust binding and neutralizing antibody responses in mice, hamsters, and nonhuman primates, resulting in robust protection against infection of both the upper and lower respiratory tracts when evaluated in the hamster model of SARS-CoV2 infection (*11*). In a recent major milestone for this platform, following the completion of a phase II/III clinical trial in India (clinical trial identifier CTRI/2021/09/036379) for the drug product HDT/Gennova COVID-19 (HGC019), emergency use approval was granted in India marking the first repRNA approved for human use. SARS-CoV-2 investigational drug products based on this platform are currently under evaluation in ongoing phase I trials in South Korea under the name QTP104, as well as in Brazil and the US under the name HDT-301. Recently, Arcturus announced (*12*) positive efficacy results in their 17,000 participant, phase III clinical trial evaluating a another repRNA-based vaccine, also referred to as self-amplifying RNA, delivered via lipid nanoparticles. In Arcturus’ trial, a 5µg prime/boost provided 55% efficacy against symptomatic disease and 95% efficacy against severe disease, similar to reports for Moderna’s 100µg prime/boost regimen when omicron and delta were the circulating variants (*13*), demonstrating the dose-sparing capacity of a repRNA vaccine approach. Here, we investigated protection afforded by repRNA-CoV2S in immunized nonhuman primates when neutralizing antibody responses were at their peak and after they had waned. We demonstrate that despite contraction of neutralizing antibody responses to very low or undetectable levels by almost 7 months post-immunization, sustained binding antibody responses and a rapid anamnestic recall response were able to mediate protection from infection and lung disease following SARS-CoV- 2 challenge. This study suggests a role for non-neutralizing binding antibody, or other adaptive immunity, in protection and provides direct evidence that protective immunological memory to COVID-19 vaccination persists and can confer protection even after neutralizing antibodies have waned to undetectable levels.

## RESULTS

### Effects of dose and interval between prime and boost on the immunogenicity of the repRNA-CoV2S vaccine

We previously developed a cationic nanocarrier, termed LION^TM^, with optimized surface chemistry designed to complex with repRNA at the nanoparticle’s surface (*11*). In contrast to lipid nanoparticle-based approaches, LION can be manufactured independently of the RNA component and, due to its enhanced stability at room temperature and at 4°C in the absence of a co-formulated RNA, can be locally stockpiled in preparation for mixing with an appropriate RNA. The latter step can be easily conducted at the pharmacy or bedside by simple inversion of a vial containing LION and RNA. This allows for simplified manufacturing of the RNA component and rapid adaptation to novel variants or future novel pandemics (*11*). We demonstrated that a single high dose of the repRNA-CoVS vaccine (250 μg) induced robust immune responses after only a single immunization. This dose was initially employed to establish safety and immunogenicity of the repRNA-CoV2S vaccine at the highest feasible dose (*14*).

Given the global crisis brought on by the COVID-19 pandemic, we employed nonhuman primates to identify the minimum dose, delivery volume, and dosage interval most likely to work in humans to support parallel clinical development activities with our repRNA-CoVS vaccine that had not been previously tested in humans. To do so, we employed an adaptive dose de-escalation strategy, staggering the enrollment of cohorts so the data generated from one cohort could be used to guide the immunization regimen (dose, interval between doses and delivery) in the next cohort.

We previously showed that priming and boosting with 50 μg doses spaced 4 weeks apart induced equivalent responses to a single 250 μg dose (*11*). Here, we compared immune responses in this 50 μg dose group to an additional group of NHPs immunized primed and boosted with 25 μg doses spaced 20 weeks apart (**Fig. 1A**). As shown in **Figure 1B**, a single 25 μg dose induced Spike-specific IgG antibody titers at 6 weeks post-immunization that were comparable to titers in the macaques that had received two 50 μg doses spaced 4 weeks apart, suggesting that antibody responses following a single dose of the repRNA-CoV2S vaccine continue to increase such that administration of a booster dose within just 4 weeks after the first dose may not provide an additional benefit. Strikingly, the 25 μg group exhibited sustained IgG titers for 20 weeks with only modest waning and a booster 25 μg dose at this time induced a significant increase in IgG responses that exceeded peak levels induced with two doses of the 50 μg dose (**Fig. 1B**). These results provide further evidence that a longer interval between the prime and booster doses may facilitate stronger immune responses by this vaccine strategy (*15*).

**Figure 1.**
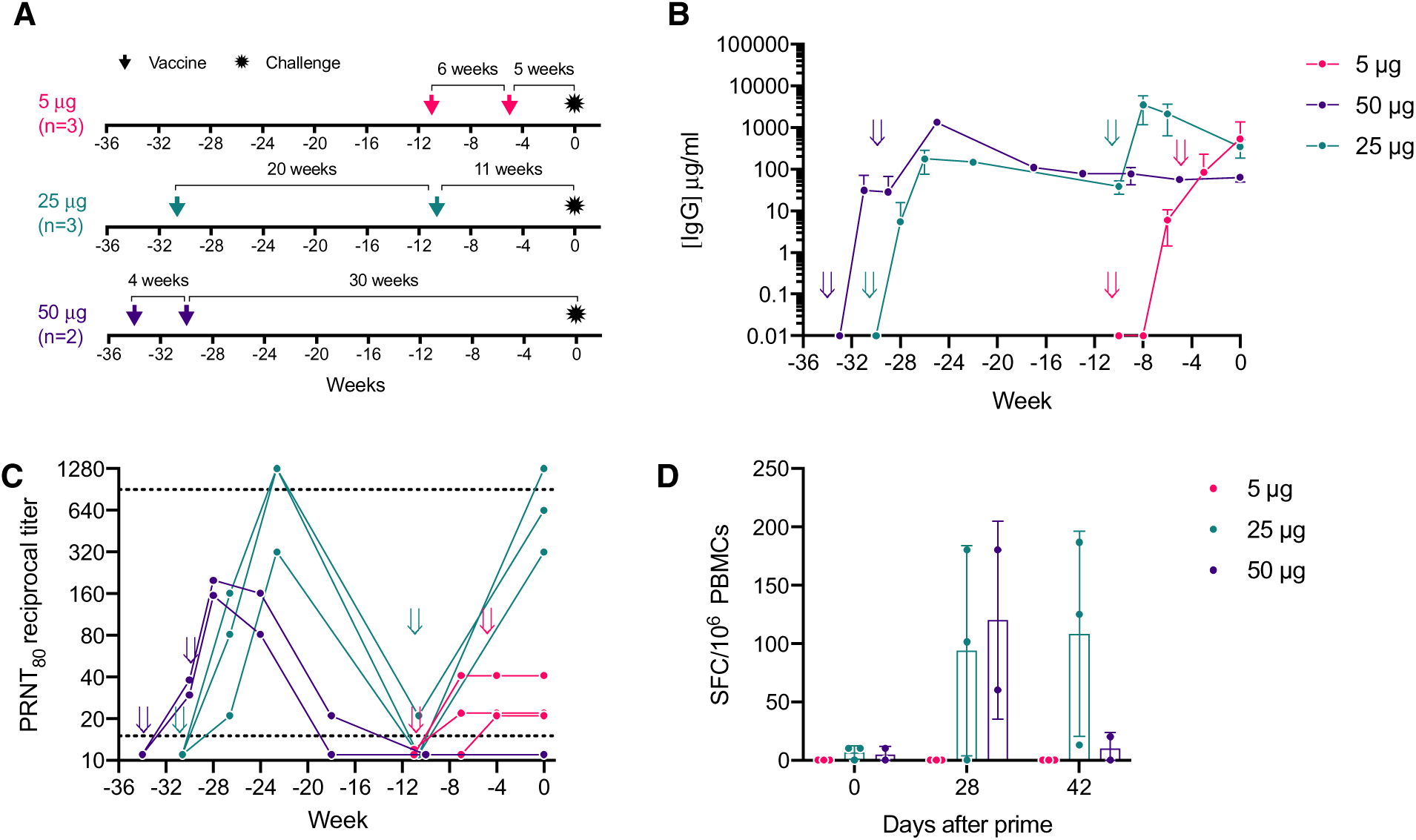
RepRNA-CoV2S induces robust and durable antibody responses in pigtail macaques. **(A)** Pigtail macaques (n=8) received two intramuscular immunizations of repRNA- CoV2S (RepRNA vax) 5 μg (n=3), 25 μg (n=3), or 50 μg (n=2) and were delivered 4-20 weeks apart as indicated in figures A-C. All animals were challenged with SARS-CoV-2 at Week 0. Blood was collected at baseline and days 10, 14, and every 14 days thereafter. **(B)** Serum anti-S IgG enzyme linked immunosorbent assays (ELISAs) and **(C)** 80% plaque-reduction neutralizing antibody titers (PRNT80) against the SARS-CoV2/WA/2020 isolate. Dotted lines indicated the lower and upper limits of detection for the assay. **(D)** Magnitude of IFN-γ T-cell analysis responses were measured by IFN-γ ELISpot assay in PBMCs following 24-hour stimulation with 11-mer overlapping peptide pools encompassing the SARS-CoV-2 spike (S) protein. All timepoints tested are post-prime except the Day 42 timepoint which corresponds to 14 days post-boost for the 50 μg group. **(B,D)** Means and standard deviations are shown.

To determine if the repRNA-CoVS vaccine could induce similar antibody responses at an even lower dose, we next investigated priming and boosting with a 5 μg dose spaced 6 weeks apart (**Fig. 1A**). IgG titers measured 2 weeks after the prime and the boost were comparable to IgG titers induced within similar timeframes in the 25 μg and 50 μg groups (**Fig. 1B**) indicating that a 5 μg dose of the repRNA/LION vaccine is sufficient to induce robust binding IgG responses in NHPs.

Neutralizing antibodies play an important role in SARS-CoV-2 viral control. We therefore measured neutralizing antibody in each group by 80% plaque-reduction neutralization test (PRNT_80_). In the 50 μg group, neutralizing antibody responses peaked 6 weeks after prime (2 weeks after the booster dose) but had waned to undetectable levels by weeks 16-24 post-prime (**Fig. 1C**). Interestingly, neutralizing antibody responses after only a single 25 μg dose peaked at 8 weeks at notably higher levels than in the group that received two 50 μg doses. By 20 weeks post-prime, neutralizing antibody responses after one 25 μg dose also declined to near undetectable levels (**Fig 1C**); however, administration of a 25 μg booster in this group restored the robust neutralizing antibody responses observed post-prime and these high levels were sustained for at least 11 weeks (the duration of the study) (**Fig. 1C**). In the 5 μg group, neutralizing antibodies peaked 4-6 weeks post-prime but the titers in all three animals were lower than peak responses induced after a single dose with the 25 μg or 50 μg doses and were not further enhanced by a booster immunization (**Fig. 1C**). Collectively, these data suggest that the 5 μg dose is suboptimal for induction of neutralizing antibody and a longer interval between prime and boost enables the development of more robust and durable binding (**Fig. 1B**) and neutralizing antibody responses (**Fig. 1C**).

Previously, we reported that doses of 50 μg or 250 μg repRNA-CoV2S induced only modest spike-specific peripheral T-cell responses that were predominantly IFN-γ secreting and directed at the receptor binding domain (*11*). Here, we evaluated if lower vaccine doses or increased spacing between vaccinations influenced the induction of SARS-CoV-2 spike-specific IFN-γ secreting T-cell responses measured by ELISPOT in peripheral blood mononuclear cells (PBMCs). Anti-spike T-cell responses measured 4 weeks after the prime immunization in the 25 μg group were of similar magnitude to levels previously reported 2 weeks post-boost in the 50 μg group (*11*) (**Fig. 1D**). Although the 5 μg dose induced robust binding antibodies on par with levels induced by the 25 μg and 50 μg doses, it did not elicit detectable T-cell responses (**Fig. 1D**). These data indicate that a 25 ug dose is sufficient to induce maximal T cell responses but a 5μg dose may be below the threshold required to induce peripheral blood T-cell responses.

### repRNA-CoV2S affords protection from viral replication in respiratory mucosa even after neutralizing antibody responses have waned to undetectable levels

With the staggered cohort enrollment, we had an opportunity to evaluate protective efficacy in animals challenged 5 (5 μg group), 11 (25 μg group) and 30 (50 μg group) weeks after the final immunization when neutralizing antibody titers were undetectable (50 μg group), low (≤ 40, 5 μg group) or high (≥ 320, 25 μg group) (**Figure 1A**). All eight vaccinated pigtails and six unvaccinated (mock) controls were challenged with SARS-CoV-2 (WA-1 strain) via combined intranasal/intratracheal routes (IN/IT). Viral loads were measured in nasal swabs collected on days 1, 3, 5 and 7, in bronchial alveolar lavages (BAL) collected on days 3, 5 and 7 and in lung tissue collected at day 7 by analysis of subgenomic RNA using qRT-PCR. Five of the 6 control animals exhibited significant viral shedding in the nasal swabs by day 3 post-infection (**Figure 2A**) and all 6 control animals exhibited high viral loads in the BAL at days 3 and 5 (**Figure 2B**) and in lung tissue at day 7 (**Figure 2C**). In contrast, by day 5 post-infection (PI), none of the 8 vaccinated animals had detectable viral RNA in the nasal swabs (**Figure 2A**) and in the BAL, 5 of 8 vaccinated animals had detectable but low viral loads (**Figure 2B**). Overall, all 3 vaccinated groups exhibited lower viral loads and accelerated viral clearance in the nasal and lung compartments compared to the controls with no significant differences between the vaccine groups. Further analysis of the area under the curve (AUC) viral loads in the nasal swabs (**Fig. 2A, right panel**) or BAL (**Fig. 2B, right panel**) between days 3 to 7 PI in all 8 vaccinated animals show significantly reduced viral loads in the vaccinated animals when compared to unvaccinated controls. A timed necropsy was performed on day 7 PI to measure viral load within the lungs, including the right-upper, middle, and lower lung lobes. As shown in **Figure 2C**, viral loads in the lung tissues of the vaccinated animals were, overall, lower than in the control group (P = 0.0006) with 4 of the 8 vaccinated animals, including all 3 animals in the 25 μg group, exhibiting no detectable virus in any of the lung tissues. Strikingly, the 50 μg group exhibited significant blunting of viral load in all specimens tested even though there was no detectable circulating neutralizing antibody in the blood at the time of challenge. Together, these data show that all 3 doses of the repRNA-CoV2s vaccine reduced and/or accelerated clearance of viral load in the upper (nasal) and lower (BAL, lung) respiratory tracts. Furthermore, these data show that the repRNACoV2s vaccine can afford sustained protection even if neutralizing antibody responses in the blood are very low (5 μg) or undetectable (50 μg group) at the time of challenge.

**Figure 2.**
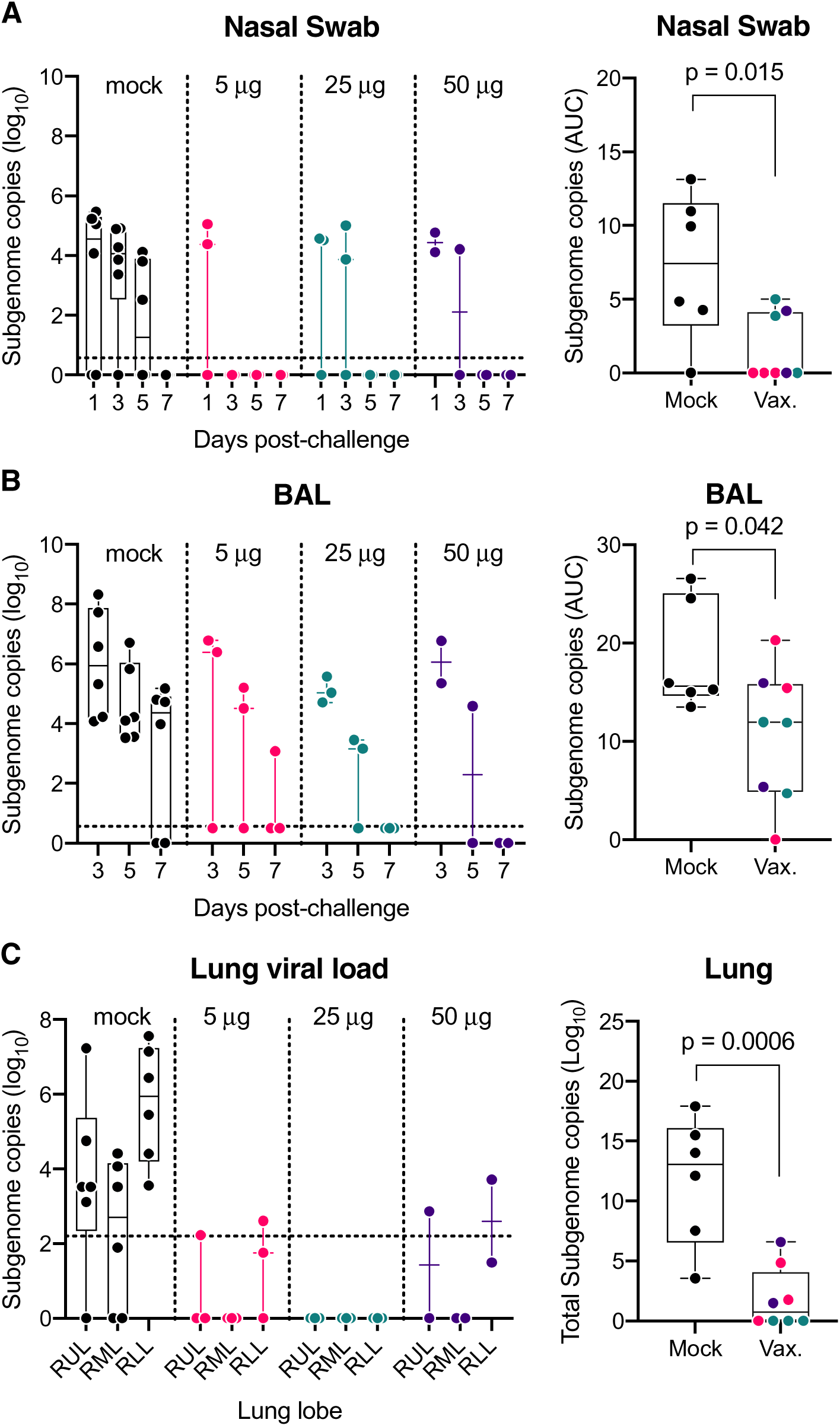
Reduced SARS-CoV-2 viral burden in respiratory mucosa in RepRNA-CoV2S vaccinated macaques. Viral loads in the indicated samples were quantified using qRT-PCR to measure subgenomic RNA. Area under the curve (AUC) viral burdens were measured by RT-PCR analysis of subgenomic RNA on days 3, 5 and 7 post infection in **(A)** nasal swabs and **(B)** bronchioalveolar lavages (BAL) and **(C)** on day 7 in the lung tissues (right lobe). RUL, right upper lung; RML, right middle lung; RLL, right lower lung. Box and Whisker plots with minimum to maximum ranges are shown. Dotted lines indicated the limit of detection for the assay. Unpaired T-test p-values are shown, with p-values ≤ 0.05 considered significant.

### The repRNA-CoV2S vaccine affords durable protection from SARS-CoV-2 induced clinical disease and pathology

We next evaluated the ability of the vaccine to protect from clinical disease and lung pathology. Animals were comprehensively evaluated by trained research staff for evidence of disease including reduced appetite, lethargy, respiratory distress, and general appearance. The most common clinical observations among unvaccinated animals were lethargy, abnormal respiration, and nasal discharge. Among the vaccinated animals, the most common observation was reduced appetite. None of the 3 animals in the 25 μg group exhibited measurable clinical scores whereas mild clinical scores occurred in 1 or 2 animals in each of the 50 μg and 5 μg groups, respectively (**Figure 3A, left panel**). When taken together, overall, clinical scores in the vaccinated animals were significantly reduced when compared to the controls with the lowest scores observed in the 25 μg group that also had the highest neutralizing antibody response at the time of challenge (**Figure 3A, right panel**).

**Figure 3.**
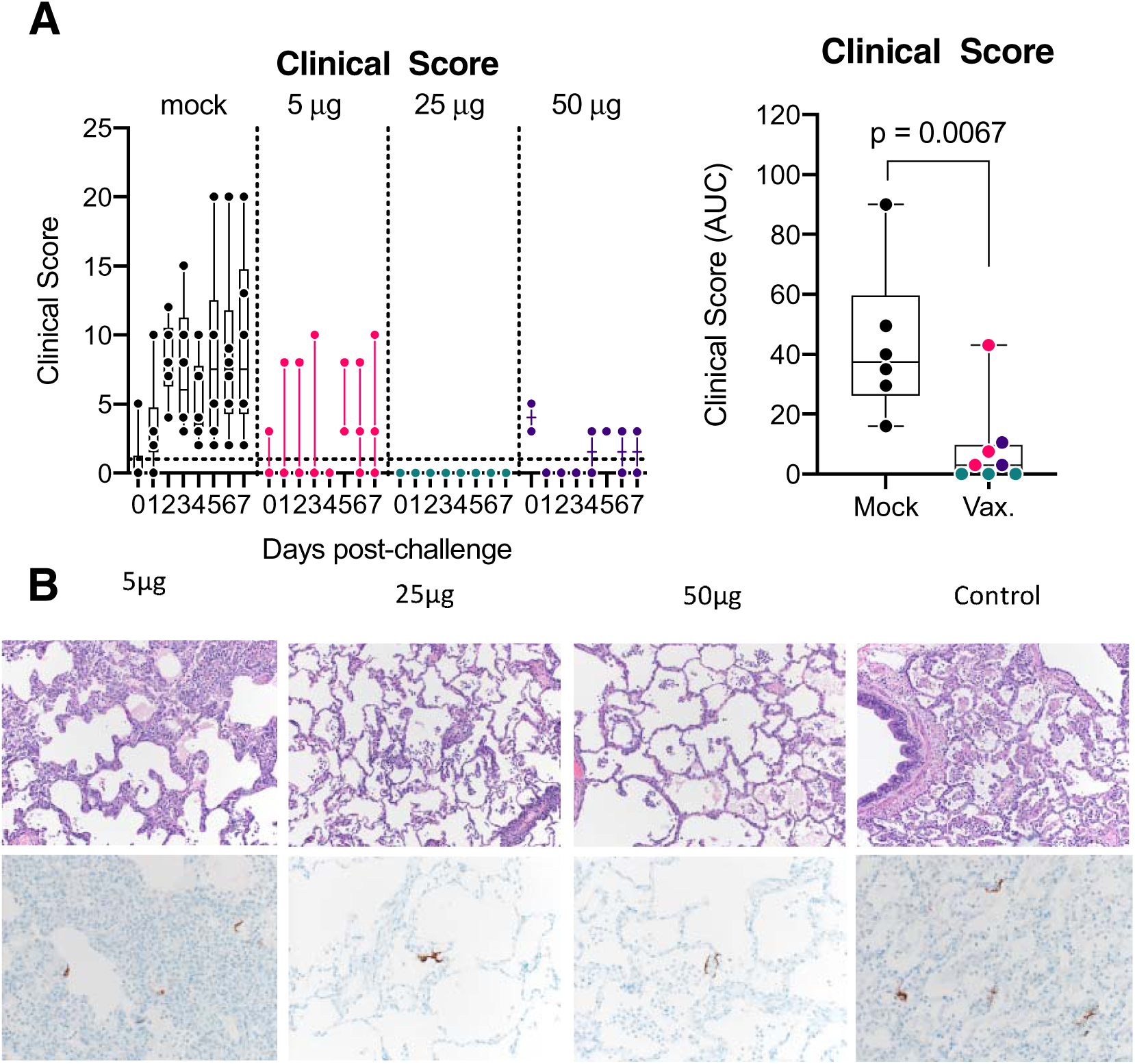
RepRNA-CoV2S vaccination reduces clinical disease and lung pathology. **(A)** Animals were comprehensively evaluated for clinical disease including activity levels, appetite, appearance, and evidence of respiratory distress. Cumulative scores are shown. Area under the curve (AUC) of clinical scores were calculated 1-7 days after infection as described in the methods. Box and Whisker plots with minimum to maximum ranges are shown. Unpaired T-test p-values are shown, with p-values ≤ 0.05 considered significant. **(B)** Formalin fixed lung tissue was sectioned and stained with H&E (top panels) for SARS-CoV-2 antigen (bottom panels) at day 7 post-challenge. All six of the monkeys in the sham vaccinated group developed some degree of pulmonary pathology when inoculated with SARS-CoV-2, predominantly in the lower lung lobes. Pathological analysis of lungs in the vaccinated animals was generally less severe than in the control animals and lesions in the 25µg were less severe overall than compared to the 5µg or 50µg groups. Rare immunoreactivity to SARS-CoV-2 nucleocapsid was seen in vaccinated groups. Representative images are shown and the complete histological findings are provided in Supplemental Table 3.

To further investigate lung pathology, on day 7 PI, formalin-fixed lung tissue was blindly evaluated for pathology. All six macaques in the unvaccinated group developed some degree of pulmonary pathology after SARS-CoV-2 challenge and predominantly in the lower lung lobes (**Figure 3C-D, Supplemental Table 2**). Lesions were characterized as multifocal, minimal to marked, interstitial pneumonia frequently centered on terminal bronchioles. The pneumonia was characterized by thickening of alveolar septa by edema fluid and fibrin and small to moderate numbers of macrophages and fewer neutrophils. There was moderate type II pneumocyte hyperplasia. Alveoli contained moderate numbers of pulmonary macrophages, neutrophils and fibrin as well as edema. Multifocally, there were perivascular infiltrates of small to moderate numbers of lymphocytes that form perivascular cuffs. Among vaccinated animals, lesions, when present, were generally less severe (**Figure 3B** **and Supplemental Table 2**) and overall, lesions in the 25 μg group, if present, were less severe compared to the 5 and 50 μg groups. The complete histological findings are provided in **Supplemental Tables 2 and 3**. Cumulatively, our data demonstrate that in addition to reducing viral burdens in the upper and lower respiratory system, all doses of the repRNA-COV2S vaccine, including the 5 and 50 μg groups that had little or no neutralizing antibody at the time of challenge, were protected against clinical disease and lung pathology 5-30 weeks after vaccination, but the most effective protection was observed in the 25 μg that had the strongest neutralizing antibody at the time of challenge.

### Vaccination reduces the cytokine response in the lung post-SARS-CoV-2 challenge

Human SARS-CoV-2 infection leads to marked increases in inflammatory cytokines that are associated with more severe COVID-19 disease (Costela-Ruiz et al., 2020). We measured cytokines and chemokines in the BAL at days 3, 5, and 7 PI (**Figure 4A, Supplemental Table 4- 6**) and blindly evaluated their association with level of clinical disease. Many pro-inflammatory cytokines including granulocyte colony stimulating factor (**G-CSF**), IFNα. MCP-1, and TGFα were positively associated with higher clinical score (**Figure 4A**). In particular, at day 5 PI higher concentrations of G-CSF, a cytokine that is associated with more severe COVID-19 clinical disease in humans (*16*) significantly correlated (p = 0.032, r = 0.578) with higher levels of clinical disease (**Fig 4A, Supplemental Table 6**).

**Figure 4.**
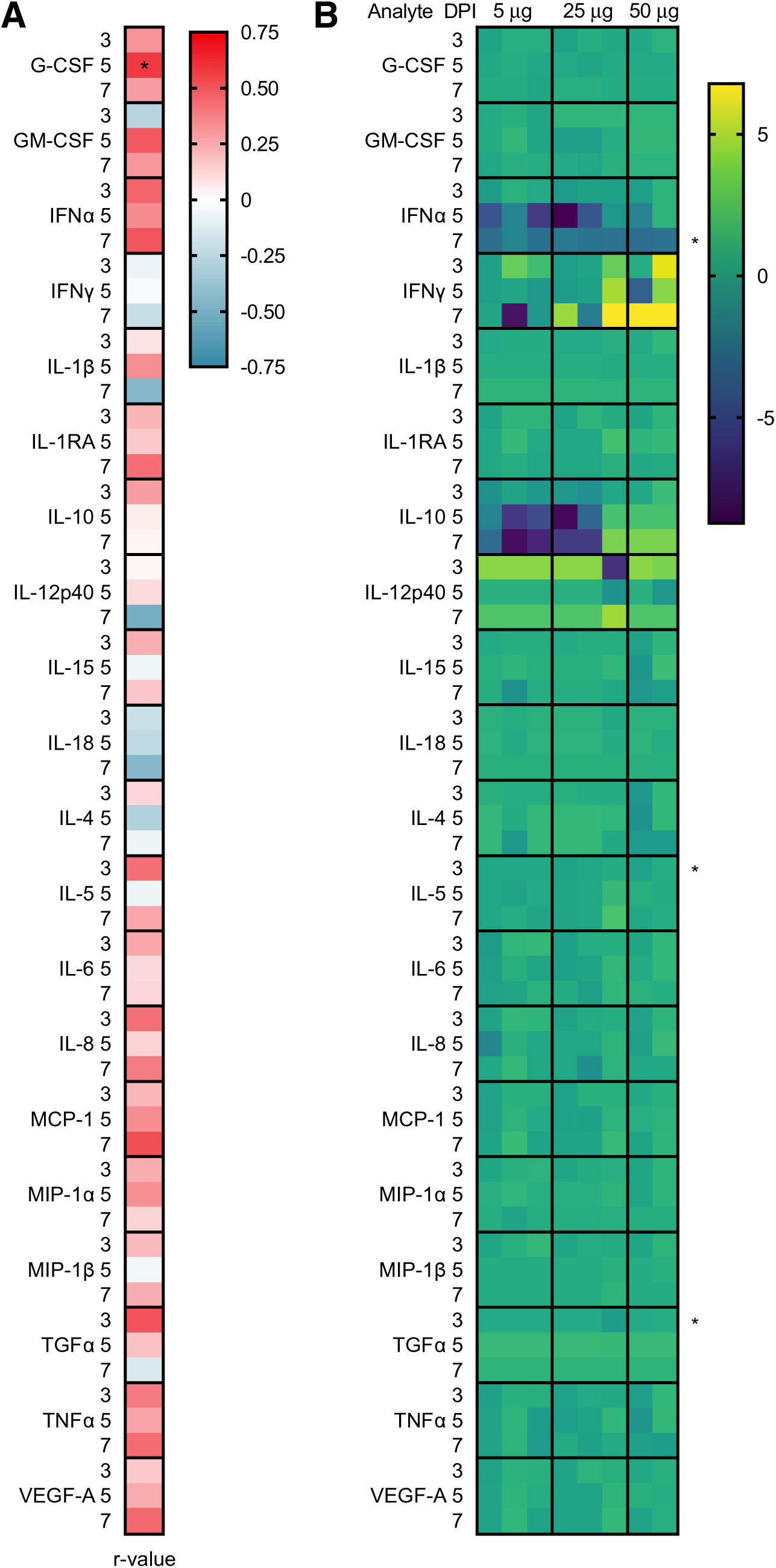
Levels of cytokine and chemokines in the BAL following SARS-CoV-2 challenge. Concentrations of cytokines and chemokines were measured in the BAL at days 3, 5 and 7 by multiplex immunoassay. **A)** BAL analytes associated with disease. Spearman rank correlation tests were performed to compare analyte concentrations in the BAL to AUC clinical scores across all animals. Shown are the r-values. A *p-value < 0.05, is considered significant. **(B)** Log transformed cytokine concentrations in the BAL of vaccinated animals were normalized to the mean responses in the mock controls, respectively, at 3, 5, and 7 days post-infection. Fold change values are displayed. A Wilcoxon test was performed to identify fold changes that are significantly different between the vaccinated and mock control groups, A *p-value < 0.05 is considered significant.

Next, to determine if the vaccine reduced inflammatory responses in the BAL, we blindly compared the levels of cytokine induction in the vaccinated animals relative to level in the control group (**Figure 4B, Supplemental Figure 1, Supplemental Table 4**). At day 3 PI, five of the six mock-treated controls had elevated levels of one or more proinflammatory cytokines that are associated with COVID-19 disease in humans (*17*) (e.g. IFNL, IL-6, IL-8, IL-10, IL-1β, TGFL, TNFL) or chemokines important for macrophage, monocyte and neutrophil recruitment (e.g. MCP-1/CCL2, MIP1L/CCL3, MIP1β/CCL4) that either declined rapidly by days 5-7 PI or persisted through day 7 PI (**Supplemental Figure 1**). In contrast, one or more of these same pro-inflammatory cytokines and chemokines were only transiently detected on days 3-5 PI in up to 5 of the 8 vaccinated animals with no apparent differences between the 3 vaccine doses (**Supplemental Figure 1**). Cytokines associated with T- and B-cell proliferation and activation (e.g. IL-4, IL-5, IL-15), indicative of a primary or recall adaptive immune response to SARS-CoV-2 infection, were also elevated in up to 5/6 control and 5/8 vaccinated animals after challenge (**Figure 4B, Supplemental Figure 1**) with a trend toward higher cytokine responses in the controls at day 3 when peak viral loads were detected in the BAL (**Fig 2B**). Collectively these data show that 5-50 μg doses of the repRNA-CoV2 vaccine administered 5-30 weeks prior to challenge effectively reduced inflammation in the lung, an outcome that is consistent with the observed lower overall clinical scores in the vaccinated group (**Figure 3A**).

### Vaccinated animals develop a robust and rapid anamnestic recall antibody response following SARS-CoV-2 challenge

All vaccinated animals showed evidence of protection from SARS-CoV-2 challenge even though 5 of the 8 vaccinated animals had low to undetectable neutralizing antibody at the time of challenge. To determine possible immune mechanisms of protection, we first investigated if a rapid post-challenge anamnestic antibody response occurred that could blunt infection at the earliest stages of the infection. The 50 μg group that was challenged 30 weeks post-immunization had undetectable neutralizing antibody at the time of challenge but exhibited a rapid and the most robust increase in both binding (**Figure 5A**) and neutralizing antibody responses (**Figure 5B**) at day 7 PI. In contrast, at the time of challenge, binding and neutralizing antibodies were highest in the 25 μg group and were only modestly boosted following SARS-CoV-2 exposure (**Figures 5A and 5B**). In the 5 μg group, despite a notable increase in post-challenge binding antibody responses (**Figure 5B**), levels of neutralizing antibody decreased to levels at or below the limit of detection, suggesting that infection in this group recalled only non-neutralizing antibody responses (**Figure 5B**). Collectively, these data demonstrate a rapid anamnestic binding and/or neutralizing antibody response occurred in all 3 vaccinated groups.

**Figure 5.**
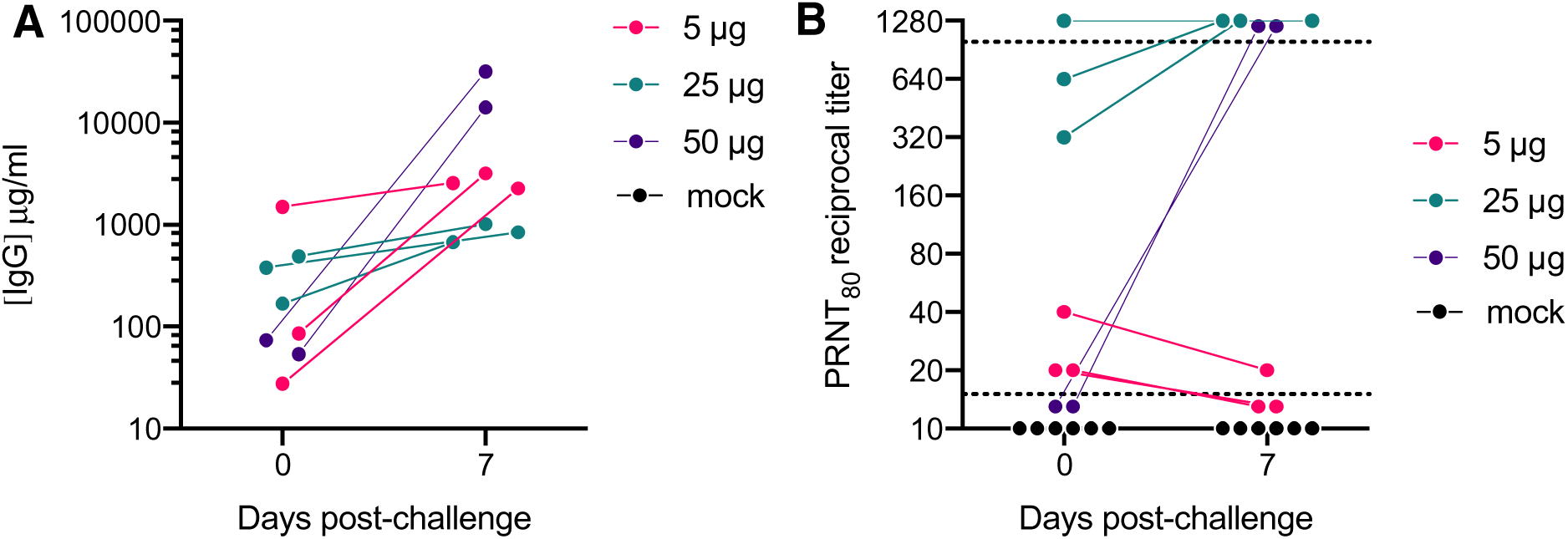
Macaques with undetectable neutralizing antibody responses prior to challenge develop a robust amnestic recall response post-challenge. Binding and neutralizing antibody responses were measured at 0 and 7 days post-challenge. Shown are (A) Serum anti- S binding antibody responses measured by IgG ELISA and (B) neutralizing antibody titers measured by PRNT80 against the SARS-CoV2/WA/2020 isolate.

Notably, 7 days post-challenge, significant viral load was still detected in the BAL and/or lung tissues of all 6 controls whereas in all 8 vaccinated animals, viral loads in the nasal swabs (**Figure 2A**) and the BAL (**Figure 2B**) had declined to undetectable levels and there were only very low levels of virus detected in the lung tissue of 4 of the 8 vaccinated animals including all 3 animals in the 25μg group that had the highest neutralizing antibody at the time of challenge (**Figure 2C**). Together, these data suggest that a rapid recall antibody response in the vaccinated animals likely contributed to protection from disease by accelerating viral clearance in the lung.

Vaccinated animals exhibited a range in viral loads in the BAL days 3-7 post-infection, enabling further analysis of immune correlates of protection. We therefore compared binding antibody, neutralizing antibody and IFN-γ secreting T-cells measured at peak levels post-vaccination, just prior to challenge and/or at 7 days post-challenge to the area under the cure (AUC) viral loads measured from days 3-7 post-infection in the BAL of the vaccinated animals (**Figure 6**). Among these comparisons, Spike binding IgG levels present prior to challenge significantly correlated with lower viral burden in the BAL (**Figure 6**) whereas neutralizing antibody and IFN-γ T-cell responses did not significantly correlate with BAL viral burden at any timepoint (**Figure 6**). Together, these data indicate a role for binding antibody responses in protection from SARS-CoV-2 following vaccination with repRNA-CoV2S/LION.

**Figure 6.**
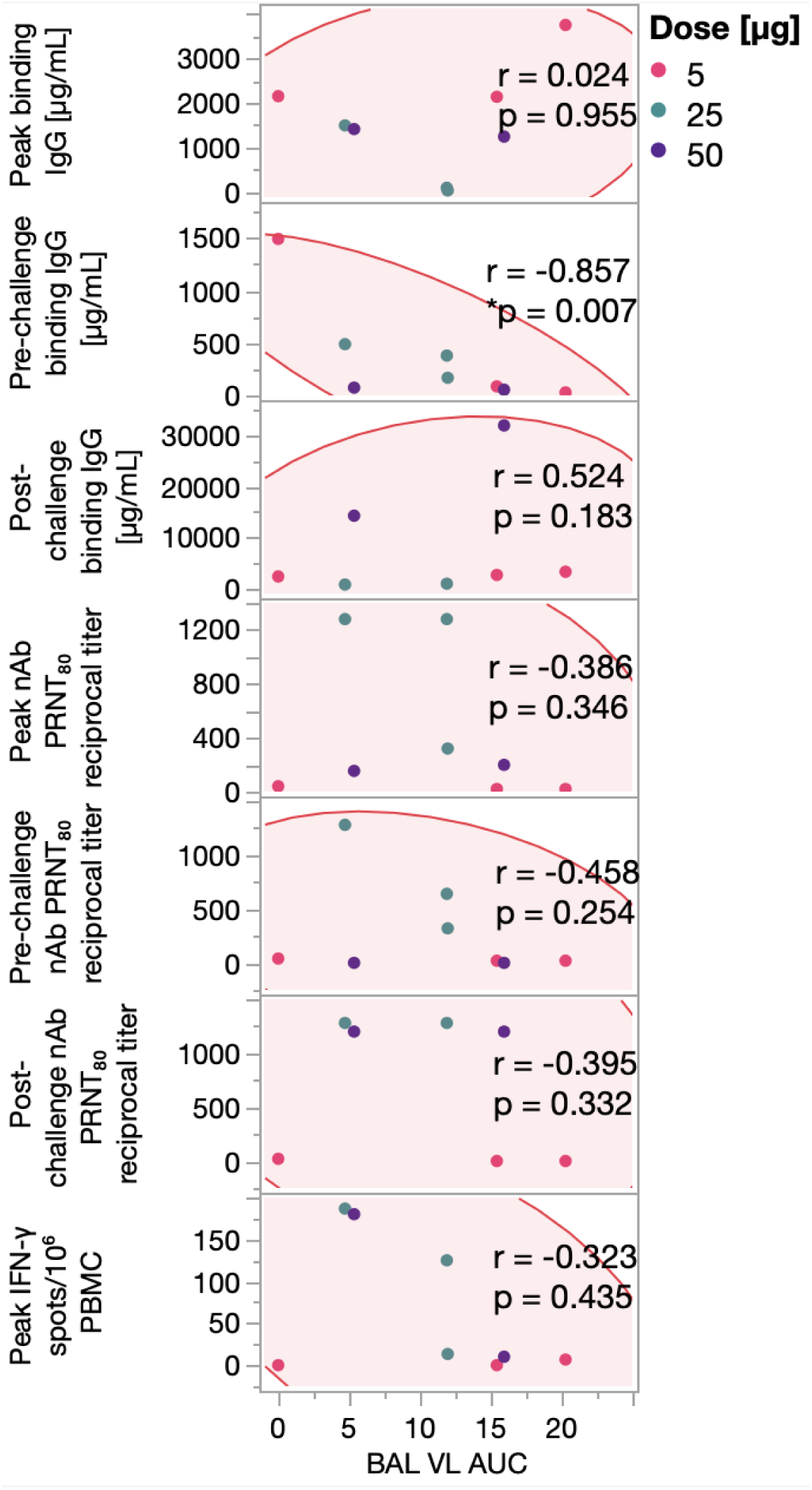
Levels of binding antibodies prior to infection correlate with reduced viral burden in BAL post-infection. Correlations between BAL viral load (VL) burden measured as area under the curve (AUC) and peak, pre-challenge and/or post-challenge (7 DPI) of Serum anti-S IgG ELISA (top panels 1-3), PRNT80 antibody titers against the SARS-CoV2/WA/2020 isolate (middle panels 4-6), and/or Magnitude of IFN-γ T-cell analysis responses measured in PBMCs following stimulation with SARS-CoV-2 spike peptides evaluated by IFN-γ ELISpot assay (bottom panel 7). Dots represent individual animals in the groups receiving 5 μg (pink), 25 μg (teal), or 50 μg (purple) repRNA-COV2 vaccines. Spearman’s rank correlation is shown, with p-values ≤ 0.05 considered significant.

## DISCUSSION

In this report we demonstrate that the repRNA-COV2S vaccine induced durable immunity against SARS-CoV-2 challenge in pigtail macaques. Vaccination with repRNA- COV2S significantly reduced viral burdens in the lungs and protected against evident disease even when animals were challenged months post-vaccination, after neutralizing antibody responses had waned to very low or undetectable levels. In addition to significantly reducing viral burden in the lungs and protecting against clinical disease, the repRNA-CoV2S vaccine also significantly reduced viral shedding in the upper airway. This is an important consideration for SARS-CoV-2 vaccines as reduction of upper respiratory tract viral loads may be important to interrupt transmission of SARS-CoV-2 from vaccinated individuals (*18*). These observations were recently recapitulated in a hamster model of variant SARS-CoV2 infections, where we demonstrated enhanced protection in the upper airway when matching repRNA-CoV2S composition to the SARS-CoV2 challenge virus (*19*). Several vaccines have been shown to protect against lower respiratory tract infection with reduced impact on viral shedding in the upper respiratory tract. In ChAdOx1 nCov-19 vaccinated rhesus macaques, significantly reduced viral burden in the lungs was not associated with reduced viral burden in the nasal swabs (*20*). Similarly, despite eliciting sterile protection in the lungs, ChAdOx1 nCov-19 vaccination did not confer reduced viral loads in the oral cavity when hamsters were challenged with the B.1.351 VoC (*21*). However, alternative vaccine platforms (*22, 23*) or intranasal rather than intramuscular administration of SARS-CoV-2 vaccines (*20, 24*) may result in significant reduction of viral shedding in the upper respiratory tract.

The repRNA-CoV2S vaccine elicited both B- and T-cell responses against SARS-CoV-2. Neutralizing antibody titers were evaluated using the stringent PRNT_80_ assay. Neutralizing antibody responses induced by this vaccine were consistent with other COVID-19 vaccine candidates tested in NHP (*25*), including the animals that received the lowest dose of 5 μg. Interestingly, in the 50 μg group, a decline in neutralization titer to undetectable levels within 5 months after the second vaccination did not correlate with a decline in ELISA titer suggesting that the waning in neutralizing antibody following vaccination was not due to a decline in overall virus-binding titers against SARS-CoV-2 spike. Similar declines in neutralizing activity but not declines in overall binding antibody titers have been observed following SARS-CoV-2 infections (*26*). However, some studies have found durable virus binding and virus neutralizing activity following either infection or vaccination (*5, 27*), whereas others have found that a decline in neutralizing titers following infection correlated with an overall decline in virus binding titers (*28*). Here, we report that despite widely different levels of neutralizing antibody from high to undetectable at the time of challenge, the vaccine afforded a significant reduction in viral burden in the respiratory mucosa, reduced clinical disease and lower lung pathology in all 3 vaccinated groups indicating the vaccine induced sufficient B-cell immunity, either circulating or memory, to mediate significant protection. Interestingly, our study revealed that higher levels of binding antibody, but not serum neutralizing antibodies, significantly correlated with reduced viral burden in the BAL. COVID-19 vaccines in NHP (*29*) and humans (*1*) have reported important roles for both neutralizing and non-neutralizing antibodies in reducing viral infection and COVID-19 disease. Further investigation into the role of non-neutralizing antibody functions in protection, including ADCC are needed (*30*).

This study demonstrates COVID-19 vaccine efficacy with no detectable levels of neutralizing antibodies at the time of challenge. The two animals in the 50 μg dose arm had reduced viral RNA in the lungs and lower clinical scores demonstrating that these animals sustained effective immune responses for at least 30 weeks after their last vaccination. The anamnestic neutralizing responses in the 25 and 50 μg groups suggest that rapid recall of neutralizing antibodies after infection may be sufficient for protection. The lack of an anamnestic nAb response observed in the 5 μg group could be due to either the lower vaccine dose or the timing between the last dose and the challenge. The significant increases in binding antibody even while neutralizing responses declined in this group, could be indicative of a skewed recall response to non-neutralizing epitopes. Prior to this study, BNT162b or mRNA- 1273 vaccination in rhesus macaques was shown to be protective 55 days (*31*) or 1 year after vaccination (*7*). However, in contrast to our findings here, detectable neutralization titers were still present in these animals prior to challenge. Data in humans demonstrating that the Pfizer and Moderna COVID-19 vaccines are effective at preventing hospitalization for at least 6 months after vaccination despite waning antibody responses (*4*) are consistent with the findings reported here. Given the observation that the presence of circulating neutralizing antibody against respiratory viruses has a tendency to wane, in contrast to durable responses against viruses that disseminate systemically (*32*), there may be too much emphasis placed on circulating neutralizing antibody as a correlate of protection from COVID-19 whereas memory responses that can rapidly recall and clear an infection may be adequate to prevent severe disease. It is possible that recall T-cell responses are also important for viral control and/or reducing immunopathology in the lung, however limited sampling post-challenge in this study precluded this analysis. Recent studies in vaccinated NHP demonstrate that failure to control Omicron in the upper respiratory tract despite neutralizing antibodies is associated with impaired CD8+ T-cell responses, suggesting a role for both antibodies and CD8 T-cells in protection (*33*).

Our study has several important limitations. First, we designed the vaccine against and used the A.1 strain of SARS-CoV-2 for the infection studies, therefore conclusions about durable protective immunity are presented in the context of homologous vaccination/infection. At the time of submission, the Omicron VoC and its related variants are the predominant circulating strains of SARS-CoV-2 in the United States (CDC) and continued evaluation of vaccine efficacy in human cohorts suggest decreased efficacy of current vaccines against VoC infection, but not necessarily disease (*36*). Although the reason for decreased protection from infection remains to be fully determined, decreased neutralization activity against VoCs (*37, 38*) and/or waning immunity (*39*) may contribute to this observation. While the evaluation of variant-specific mRNA vaccines is ongoing by us and others, it appears that variant-specific vaccines may offer the most advantage in the context of naïve individuals (*19*) as Moderna recently demonstrated that a B.1.351-specific vaccine booster in A.1-immune individuals did not improve neutralizing antibody responses against B.1.351 virus compared to those who received an A.1 booster (*40*). We evaluated this same vaccine candidate against different VoC infections in the hamster model including B.1.351 and B.1.1.7 and demonstrated substantial protection against each of these variants (*19*). Second, the interpretation of the BAL cytokine data is limited by the lack of a baseline sampling; thus, we cannot exclude the possibility that cytokine levels prior to vaccination may have influenced these results. Furthermore, it is possible that the BAL collection procedure induced a cytokine response, which reduces our ability to interpret cytokines induced by the infection versus procedures. Third, we were unable to assay for infectious virus in respiratory tract samples due to depletion of the required sample volumes, however in our evaluation of protective efficacy in the hamster model, we found that the subgenomic N assay was more sensitive than the infectious viral load assay adding more stringency to our statistical analyses. Lastly, although we evaluated different doses and timings between vaccinations, we did not power or design the study to identify the most optimal vaccine regimen, however, these questions are being directly addressed in ongoing studies and clinical trials.

Cumulatively, our data demonstrate that the repRNA-CoV2S vaccine elicits protective and durable immunity against SARS-CoV-2 in a non-human primate model of disease. Vaccination significantly reduced viral replication and virus-induced pathology in the lungs, improved clinical outcome and reduced viral shedding. These findings indicate that the repRNA- CoV2S vaccine may not only protect against severe COVID-19 but also interrupt transmission of SARS-CoV-2. Furthermore, this study highlights an important role for memory B cell responses in mediating durable protection from disease even after neutralizing antibody responses have waned. These data support continued development of this vaccine in pre-clinical and clinical studies.

## MATERIALS AND METHODS

### Biosafety and Ethics

All procedures with infectious SARS-CoV2 were conducted at high biocontainment conditions in accordance with operating procedures approved by the Rocky Mountain Laboratories institutional biosafety committee. Animal experiments were approved by the institutional animal care and use committee and performed by experienced personnel under veterinary oversight. Sample inactivation prior to analyses followed IBC approved protocols(*41*).

### Pigtail Macaques and Sample Collection

Fourteen male pigtail macaques (*Macaca nemestrina*) were utilized in this study, see **Supplemental Table 1** for animal characteristics. Two animals previously received irrelevant vaccinations with a combination of Hepatitis B virus (HBV) DNA and protein vaccines consisting of HBV core and surface antigens conjugated to anti-CD180 to promote antigen targeting to B cells (**Supplemental Table 1**). All animals were initially housed at the Washington National Primate Research Center (WaNPRC) for the COVID-19 vaccination phase and then were transferred to Rocky Mountain Laboratories (RML) for the SARS-CoV-2 challenge phase. Both facilities are accredited by the American Association for the Accreditation of Laboratory Animal Care International (AAALAC), and all procedures performed on the animals were with the approval of the University of Washington and RML’s Institutional Animal Care and Use Committee (IACUC). Animals were housed in individual cages with 12h light-dark cycles. Animals were provided with commercial monkey chow, treats and fruit by trained personnel. Water was available ad libitum. They were offered environmental enrichment including a variation of toys, videos, music and human interaction.

### RepRNA-CoV2S Vaccination

The vaccine was prepared as previously described (*11*) using a Lipid InOrganic Nanoparticle (LION) complexed with an alphavirus-derived replicating RNA (repRNA) encoding the full-length spike of SARS-CoV2 (repRNA-CoV2S). The vaccines were delivered by i.m. injection into the quadriceps and deltoid muscles over 5 sites. Animals received 2 doses of either a 5 μg (n=3), 25 μg (n=3), or 50 μg (n=2) dose of repRNA-CoV2S or remained unvaccinated (mock). The timeframe between vaccine doses was as follows: 4 weeks (50 μg), 6 weeks (5 μg), or 20 weeks (25 μg). The vaccine was delivered in a total volume 250-1250 μl over 1-5 sites. These conditions were employed as we were simultaneously evaluating the safety and immunogenicity of different administration techniques, dosages and intervals between doses to support the development of a pharmacy manual to be included in parallel clinical development activities. (see **Supplementary Table 1** for details). All injection sites were monitored post-immunization for any signs of local reactogenicity, with no adverse events reported.

### SARS-CoV-2 Viral Challenge

SARS-CoV-2 strain nCoV-WA1-2020 (MN985325.1) was provided by CDC, Atlanta, USA. Virus propagation, titration and sequence confirmation of identity was performed as previously described (*20, 42*). Virus propagation was performed with a single passage in VeroE6 cells in DMEM supplemented with 2% fetal bovine serum, L-glutamine and penicillin and streptomycin. Virus was titered by median tissue culture infectious dose (TCID_50_) on VeroE6 cells and was sequenced to confirm identity and exclude contamination.

### IFN-γ Enzyme-linked immunospot assay (ELISPOT)

Antigen-specific T-cells secreting IFN-γ in the PBMCs were detected using a Human IFN-γ Single-Color ELISPOT or Human IL-4/IFN-γ Double-Color ELISPOT (ImmunoSpot, Shaker Heights, Cleveland, OH), per the manufacturer’s protocol. Briefly, cryopreserved PBMC cells were thawed, and 1-3 x 10^5^ cells were stimulated for 24-48 hours with 11 SARS-CoV-2 Spike peptide pools (17- or 18-mers with 11 amino acid overlap) (Genscript, Piscataway, NJ) at a concentration of 1μg/mL per peptide. DMSO and Concanavalin A (ThermoFisher, Waltham, MA) were used as negative and positive controls, respectively, as previously described (Erasmus et al., 2020). Spots were counted on an Immunospot Analyzer with CTL Immunospot Profession Software (Cellular Technology Ltd., Shaker Heights, Cleveland, OH). Spot forming cells (SFC) were computed following DMSO subtraction and were considered positive if the number of SFC was > 10 SFC per 1 x 10^6^ cells.

### SARS-CoV-2 Binding ELISA

Antigen-specific IgG responses were detected in untreated sera (pre-challenge) or gamma irradiated (2 megarads) sera (post-challenge) by enzyme linked immunosorbent assay (ELISA), and performed as previously described (*11*). Briefly, ELISA plates were coated with 1 μg/ml recombinant SARS-CoV-2 S protein (*43*) and serially diluted serum samples were added and detected via anti-monkey IgG-HRP (Southern Biotech, Birmingham, AL). Plates were developed using a TMB substrate (source) and were analyzed at 405nm (ELX808, Bio-Tek Instruments Inc). IgG serum concentrations were determined from a standard curve, as previously described (*11*).

### SARS-CoV-2 Neutralization Assay

Antibody neutralization against SARS-CoV2/WA/2020 (BEI Resources, Manassas, VA) was determined by eighty percent (80%) plaque-reduction neutralizing antibody titers (PRNT_80_) in three-fold serial diluted serum, as previously described (*11*). Briefly, serum and virus were incubated for 30 min at 37°C on Vero E6 cells (ATCC, Manassas, VA). Following adsorption, plates were washed and a 0.2% agarose in DMEM supplemented with 1% FBS was overlaid onto the cells and incubated for 2 days at 37°C. Following fixation with 10% formaldehyde (Sigma-Aldrich, St. Louis, MO), cells were stained with 1% crystal violet (Sigma-Aldrich, St. Louis, MO) in 20% EtOH (Fisher Scientific, Waltham, MA). Plaques were enumerated and percent neutralization was calculated relative to the virus-only control.

### Quantitative PCR

RNA from tissues was extracted using RNeasy extraction kit (Qiagen) and RNA from BAL or nasal swabs was extracted using Qiamp extraction kit (Qiagen). SARS-CoV2 subgenomic viral RNA was quantified using primer probe sets as previously described (Wölfel et al., 2020) and Quantifast One-Step RT-PCR master mix (Qiagen) on a QuantStudio 3 or 5 instrument (ThermoFisher). A standard curve of viral RNA of known copy number was run in parallel.

### Histology and Immunohistochemistry

Tissues were fixed in 10% Neutral Buffered Formalin x2 changes, for a minimum of 7 days. Tissues were placed in cassettes and processed with a Sakura VIP-6 Tissue Tek, on a 12-hour automated schedule, using a graded series of ethanol, xylene, and PureAffin. Embedded tissues are sectioned at 5um and dried overnight at 42 degrees C prior to staining. Specific anti-CoV immunoreactivity was detected using GenScript U864YFA140-4/CB2093 NP-1 at a 1:1000 dilution. The secondary antibody is the Vector Laboratories ImPress VR anti-rabbit IgG polymer cat# MP-6401. The tissues were then processed for immunohistochemistry using the Discovery Ultra automated strainer (Ventana Medical Systems) with a ChromoMap DAB kit Roche Tissue Diagnostics cat#760-159. Slides were scored for pathology by pathologists blinded to study groups. The histological findings in the unvaccinated animals have also been reported in a comparative review of non-human primate models for SARS-CoV-2 (*44*).

### Multiplex immunoassay

Cytokine and chemokine levels in gamma irradiated (2 megarads) BAL specimens were analyzed using a custom ProcartaPlex immunoassay (ThermoFisher Scientific) nonhuman primate cytokine magnetic bead panel premixed 24-plex kit, per the manufacturer’s protocol. The levels of the analytes were assessed on a Bio-Plex 200 system (Bio-Rad).

### Statistics

Indicated statistical comparisons as described in the figure legends were performed using Prism v9 (Graphpad). Disease outcomes between mock and vaccinated animals were determined by unpaired T-test.

## List of Supplementary Materials

Fig. S1

Table S1 to S6

## Acknowledgments

We thank the WaNPRC and RML animal staff for the excellent care of the animals.

## Author contributions

Conceptualization: MAO, DWH, HF, DHF, JHE

Methodology: KMW, SL, WS, SM, JA, TBL, BB, NI, CA, SW, WG, KAG, PT, JL, GS, PTE

Visualization: MAO, DWH, WS, PTE, AK, JHE Funding acquisition: DHF, JHE, HF

Project administration: DHF, JHE, HF Supervision: DHF, JHE, MAO, HF

Writing – original draft: MAO, DWH, HF, DHF, JHE

Writing – review & editing: MAO, DWH, KMW, SL, WS, SM, JA, TBL, BB, NI, CA, SW, WG, KAG, PT, JL, GS, PTE, AK, HF, DHF, JHE

## Data and materials availability

All data are available in the main text or the supplementary materials.

---

**Supplemental Figure 1.**
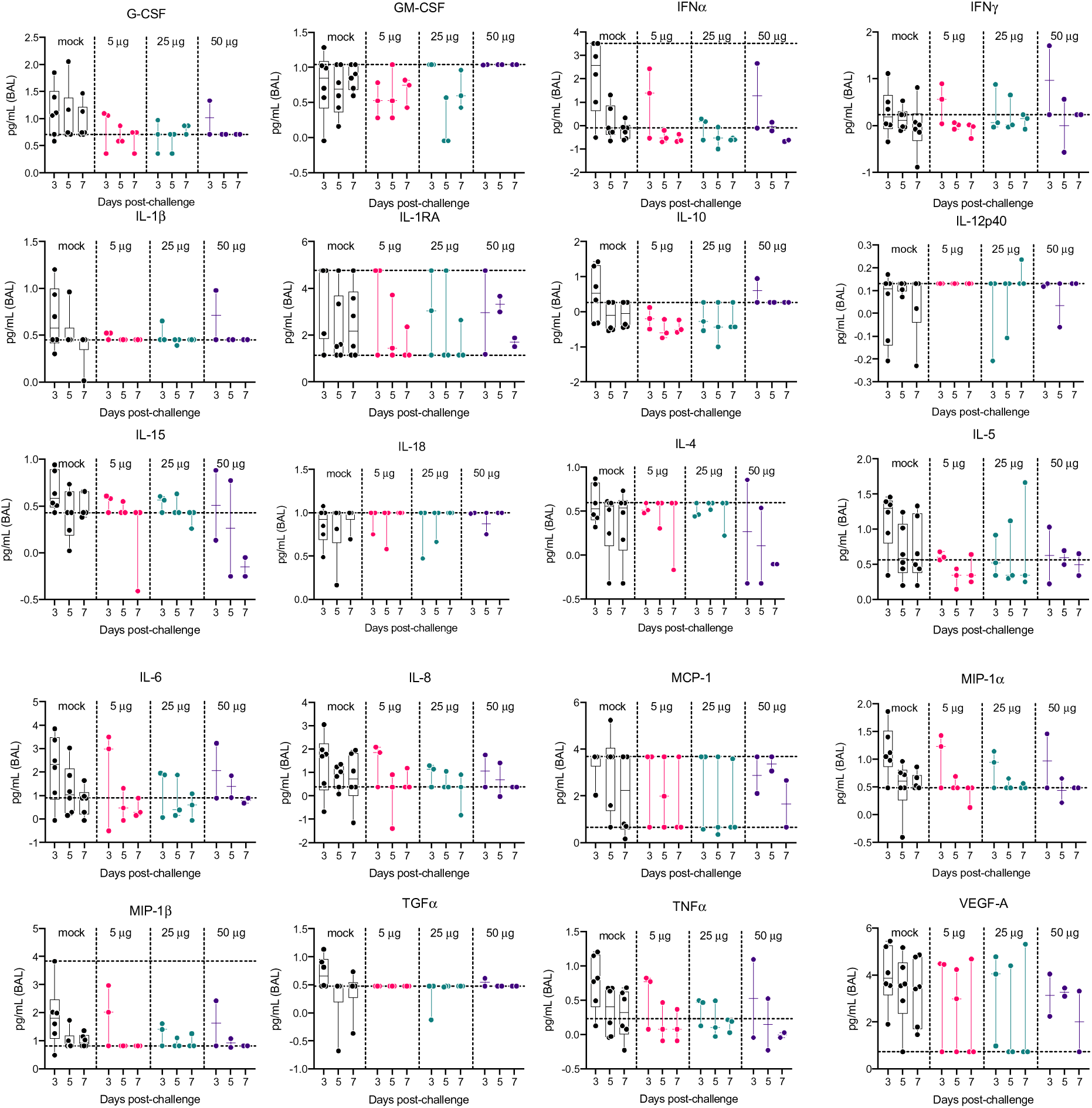
Concentration of BAL cytokines and chemokines. Log transformed concentrations of cytokines and chemokines measured in BAL by multiplex immunoassay. Horizontal dotted lines represent the lower (LLOQ) and upper (ULOQ) limits of quantification for the assay.

## References

1. P. B. Gilbert et al., Immune correlates analysis of the mRNA-1273 COVID-19 vaccine efficacy clinical trial. Science 375, 43–50 (2022).

2. K. McMahan et al., Correlates of protection against SARS-CoV-2 in rhesus macaques. Nature 590, 630–634 (2021).

3. X. He et al., Low-dose Ad26.COV2.S protection against SARS-CoV-2 challenge in rhesus macaques. Cell 184, 3467–3473 e3411 (2021).

4. N. Doria-Rose et al., Antibody Persistence through 6 Months after the Second Dose of mRNA-1273 Vaccine for Covid-19. N Engl J Med 384, 2259–2261 (2021).

5. A. T. Widge et al., Durability of Responses after SARS-CoV-2 mRNA-1273 Vaccination. N Engl J Med 384, 80–82 (2021).

6. F. P. Polack et al., Safety and Efficacy of the BNT162b2 mRNA Covid-19 Vaccine. N Engl J Med 383, 2603–2615 (2020).

7. M. Gagne et al., Protection from SARS-CoV-2 Delta one year after mRNA-1273 vaccination in rhesus macaques coincides with anamnestic antibody response in the lung. Cell 185, 113–130 e115 (2022).

8. M. Shrotri et al., Spike-antibody waning after second dose of BNT162b2 or ChAdOx1. The Lancet 398, 385–387 (2021).

9. T. Brosh-Nissimov et al., BNT162b2 vaccine breakthrough: clinical characteristics of 152 fully vaccinated hospitalized COVID-19 patients in Israel. Clin Microbiol Infect 27, 1652–1657 (2021).

10. C. Kuhlmann et al., Breakthrough infections with SARS-CoV-2 omicron despite mRNA vaccine booster dose. The Lancet 399, 625–626 (2022).

11. J. H. Erasmus et al., An Alphavirus-derived replicon RNA vaccine induces SARS-CoV-2 neutralizing antibody and T cell responses in mice and nonhuman primates. Sci Transl Med 12, (2020).

12. . (2022).

13. H. F. Tseng et al., Effectiveness of mRNA-1273 against SARS-CoV-2 Omicron and Delta variants. Nat Med, (2022).

14. J. H. Erasmus et al., Intramuscular Delivery of Replicon RNA Encoding ZIKV-117 Human Monoclonal Antibody Protects against Zika Virus Infection. Mol Ther Methods Clin Dev 18, 402–414 (2020).

15. A. P. S. Munro et al., Safety and immunogenicity of seven COVID-19 vaccines as a third dose (booster) following two doses of ChAdOx1 nCov-19 or BNT162b2 in the UK (COV-BOOST): a blinded, multicentre, randomised, controlled, phase 2 trial. Lancet 398, 2258–2276 (2021).

16. J. Guo et al., Cytokine Signature Associated With Disease Severity in COVID-19. Front Immunol 12, 681516 (2021).

17. M. Buszko et al., The dynamic changes in cytokine responses in COVID-19: a snapshot of the current state of knowledge. Nat Immunol 21, 1146–1151 (2020).

18. Y. Wu et al., A recombinant spike protein subunit vaccine confers protective immunity against SARS-CoV-2 infection and transmission in hamsters. Sci Transl Med 13, (2021).

19. D. W. Hawman et al., SARS-CoV2 variant-specific replicating RNA vaccines protect from disease following challenge with heterologous variants of concern. Elife 11, (2022).

20. N. van Doremalen et al., ChAdOx1 nCoV-19 vaccine prevents SARS-CoV-2 pneumonia in rhesus macaques. Nature 586, 578–582 (2020).

21. R. J. Fischer et al., ChAdOx1 nCoV-19 (AZD1222) protects Syrian hamsters against SARS-CoV-2 B.1.351 and B.1.1.7. Nat Commun 12, 5868 (2021).

22. K. S. Corbett et al., SARS-CoV-2 mRNA vaccine design enabled by prototype pathogen preparedness. Nature 586, 567–571 (2020).

23. N. B. Mercado et al., Single-shot Ad26 vaccine protects against SARS-CoV-2 in rhesus macaques. Nature 586, 583–588 (2020).

24. A. O. Hassan et al., A Single-Dose Intranasal ChAd Vaccine Protects Upper and Lower Respiratory Tracts against SARS-CoV-2. Cell 183, 169–184 e113 (2020).

25. P. J. Klasse, D. F. Nixon, J. P. Moore, Immunogenicity of clinically relevant SARS-CoV- 2 vaccines in nonhuman primates and humans. Sci Adv 7, (2021).

26. S. Marot et al., Rapid decline of neutralizing antibodies against SARS-CoV-2 among infected healthcare workers. Nat Commun 12, 844 (2021).

27. A. Wajnberg et al., Robust neutralizing antibodies to SARS-CoV-2 infection persist for months. Science 370, 1227–1230 (2020).

28. K. H. D. Crawford et al., Dynamics of Neutralizing Antibody Titers in the Months After Severe Acute Respiratory Syndrome Coronavirus 2 Infection. J Infect Dis 223, 197–205 (2021).

29. K. S. Corbett et al., Immune correlates of protection by mRNA-1273 vaccine against SARS-CoV-2 in nonhuman primates. Science 373, eabj0299 (2021).

30. G. J. Rieke et al., Induction of NK cell-mediated antibody-dependent cellular cytotoxicity (ADCC) against SARS-CoV-2 after natural infection is more potent than after vaccination. J Infect Dis, (2022).

31. A. B. Vogel et al., BNT162b vaccines protect rhesus macaques from SARS-CoV-2. Nature 592, 283–289 (2021).

32. J. W. Yewdell, Individuals cannot rely on COVID-19 herd immunity: Durable immunity to viral disease is limited to viruses with obligate viremic spread. PLoS Pathog 17, e1009509 (2021).

33. A. Chandrashekar et al., Vaccine protection against the SARS-CoV-2 Omicron variant in macaques. Cell, (2022).

34. M. C. Chang, S. Hild, F. Grieder, Nonhuman primate models for SARS-CoV-2 research: Consider alternatives to macaques. Lab Anim (NY) 50, 113–114 (2021).

35. A. Melton et al., The pigtail macaque (Macaca nemestrina) model of COVID-19 reproduces diverse clinical outcomes and reveals new and complex signatures of disease. PLoS Pathog 17, e1010162 (2021).

36. J. Lopez Bernal et al., Effectiveness of Covid-19 Vaccines against the B.1.617.2 (Delta) Variant. N Engl J Med 385, 585–594 (2021).

37. D. Planas et al., Reduced sensitivity of SARS-CoV-2 variant Delta to antibody neutralization. Nature 596, 276–280 (2021).

38. V. V. Edara et al., Infection and Vaccine-Induced Neutralizing-Antibody Responses to the SARS-CoV-2 B.1.617 Variants. N Engl J Med 385, 664–666 (2021).

39. A. Fowlkes et al., Effectiveness of COVID-19 Vaccines in Preventing SARS-CoV-2 Infection Among Frontline Workers Before and During B.1.617.2 (Delta) Variant Predominance - Eight U.S. Locations, December 2020-August 2021. MMWR Morb Mortal Wkly Rep 70, 1167–1169 (2021).

40. A. Choi et al., Safety and immunogenicity of SARS-CoV-2 variant mRNA vaccine boosters in healthy adults: an interim analysis. Nat Med 27, 2025–2031 (2021).

41. E. Haddock, F. Feldmann, W. L. Shupert, H. Feldmann, Inactivation of SARS-CoV-2 Laboratory Specimens. Am J Trop Med Hyg 104, 2195–2198 (2021).

42. K. Rosenke et al., Defining the Syrian hamster as a highly susceptible preclinical model for SARS-CoV-2 infection. Emerg Microbes Infect 9, 2673–2684 (2020).

43. A. C. Walls et al., Structure, Function, and Antigenicity of the SARS-CoV-2 Spike Glycoprotein. Cell 181, 281–292 e286 (2020).

44. C. S. Clancy et al., Histologic pulmonary lesions of SARS-CoV-2 in 4 nonhuman primate species: An institutional comparative review. Vet Pathol, 3009858211067468 (2021).

